# A Novel Precision-cut lung slice stretch model using removable inflation materials

**DOI:** 10.64898/2026.02.01.703167

**Authors:** Cassidy Potter, Jeannie Haak, David Dean, Andrew M. Dylag, Jared A. Mereness

## Abstract

Stretch is an important biomechanical stimulus facilitating tissue development in the respiratory system by programming the epithelium, endothelium, and extracellular matrix (ECM). Lung tissue undergoes stretch induced lung differentiation under normal prenatal and postnatal development. Furthermore, supraphysiological and aberrant stretch responses are known mechanisms of acute lung injury and ECM disruption. Current *in vitro* human tissue cyclic mechanical stretch (CMS) models suffer from significant, well-recognized disadvantages and are poorly validated *in vivo* for longer-term study. *In vitro* precision-cut lung slice (PCLS) models are commonly used to study the complex structural arrangement and cellular interactions of human tissue, as well as various lung diseases, including BPD.^3^ PCLS maintain lung tissue architecture and the variety of cell types present in the lung, allowing for a more realistic imitation of the lung microenvironment.^3^ Existing agarose-inflated PCLS models are hindered by retention of agarose media in the tissue, affecting material properties and complicating stretch studies. Our novel PCLS approach utilizes several technical innovations including a removable hydrogel for inflation and uses supportive poly(ethylene glycol) (PEG) hydrogels enable improved viability and phenotype retention during cyclic mechanical stretch (CMS). This platform will induce PCLS CMS for biochemical assays (e.g. transcriptomics, proteomics) after exposure.

## 1.0 Introduction

Precision-cut lung slices (PCLS) are a powerful ex vivo model consisting of uniform 150-500 μm sections of lung tissue that preserves the complex three-dimensional architecture and multicellular composition of the lung, allowing investigators to study epithelial, endothelial, smooth muscle, and resident immune cell interactions within intact airways and parenchyma.^1^ Because PCLS maintain physiological responses, they enable direct assessment of airway contraction, calcium signaling, mediator release, and inflammatory responses while reducing reliance on in vivo models.^2, 3^ Further, PCLS can be generated from human lung tissue, thereby enhancing their translational relevance for examining respiratory infections, exposures, and disease, such as bronchopulmonary dysplasia (BPD), asthma, COPD, fibrosis, and viral infection, and for evaluating drug responses in patient-derived samples.^1, 4^ PCLS also support longitudinal studies and parallel testing of multiple conditions from a single lung, improving experimental efficiency and reproducibility. In addition, PCLS support toxicology and environmental exposure studies, enabling assessment of particle, pollutant, and cigarette smoke effects on airway and alveolar regions within the same slice.^5^ More recently, PCLS have been adopted for mechanobiology and wound-healing studies, where stretch, matrix stiffness, and epithelial repair can be evaluated under defined conditions.^6, 7^ These advantages make PCLS offer significant improvement for many types of studies over isolated cell culture or whole-animal models for mechanistic and pharmacological research.^1, 2, 4^

Clinical studies consistently show that reduced exposure to mechanical ventilation is associated with better pulmonary outcomes^8-10^. Despite these findings, the contributions of individual exposures (i.e., viral, mechanical ventilation, hyperoxia) are poorly understood because the exposures happen simultaneously and there is no suitable non-invasive way to sample human lung tissue. PCLS are an appealing model, and have also used in cyclic stretch studies^11-18^. However, PCLS culture systems are hindered by short culture viability (5-7 days), tissue necrosis, gene expression changes over time, and fibroblast proliferation.^19, 20^ Despite this, a few studies have shown the potential for these cultures to be kept beyond 15 days, but these models still face challenges^19, 21^. For example, the retention of agarose inflation media, although necessary for tissue slicing, results in a stiff hydrogel material throughout the airspaces, thereby impacting the relaxation and mechanical properties of the tissue and potentially leading to tissue stretching to supraphysiological levels in CMS studies.

Precision-cut lung slices (PCLS) require inflation with low-melting-point agarose to provide the structural support necessary for vibratome sectioning and to prevent collapse of the delicate alveolar and airway architecture. However, this process introduces several methodological complications that can affect downstream applications. The large volume of agarose retained within PCLS can impede efficient RNA extraction, often resulting in low-yield or low-quality RNA unless specialized purification strategies are used.^22^ Additionally, the presence of agarose can alter tissue stiffness and potentially obscure alveolar and airway epithelial cell surfaces, antigenic epitopes alter tissue mechanical properties.^23^ However, this prevents the tissue from retracting to a relaxed state, which could significantly impact the physiological relevance and accuracy of tissue elongation measurements in cyclic stretch studies. Finally, achieving consistent agarose temperature and delivery during lung inflation is technically demanding—agarose that gels too quickly can produce uneven stiffness and microtears, while excessive pressure during instillation can mechanically damage delicate lung parenchyma, contributing further to slice variability and reduced viability.^24^ Agarose inflation can impede downstream molecular assays such as RNA extraction, and the slicing process can result in uneven or damaged slices.^22^ For these reasons, agarose is commonly held at 60 **°**C before inflation, which can be damaging to tissue and may cause

Removing agarose from PCLS offers several benefits that enhance their utility in research and experimental applications. Agarose removal can restore the tissue’s native mechanical properties, allowing for more accurate assessments of lung mechanics and physiological responses to mechanical stimuli.^25^ This may also improve the diffusion kinetics of drugs and biochemical molecules within the tissue, thereby facilitating more reliable pharmacological and toxicological studies. Furthermore, eliminating agarose can improve the recovery of biochemical components (RNA, DNA, protein), enhancing the precision and reproducibility of biochemical assays.^26^ In the following studies, we describe an improved lung inflation method utilizing ultra-low-melting-point (ULMP) agarose that can be removed from generated PCLS. Further, we integrate these agarose-free PCLS with PEG hydrogels to permit the application of cyclic mechanical stretch without gluing or clamping, reducing tissue damage and significantly increasing the potential duration of mechanical stretch experiments over current methods.

## 2.0 Methods

### 2.1 Agarose Gel Mechanical Testing

Molten agarose solution was pipetted in 40 μl volumes into cylindrical molds with 5 mm diameters. Samples were prepared of low-melt point (LMP) agarose (16520050, Thermo Fisher, Waltham, MA, USA) and ULMP agarose (A2576, Sigma Aldrich, St. Louis, MO, USA) at concentrations of 1%, 1.5%, 2%, 2.5%, and 3% w/v, in complete DMEM media (11995065, Gibco) containing 10% FBS (SH3007103, Cytiva, Marlborough, MA, USA), and 1% antibiotic/antimycotic (15240062, Gibco) with 3 replicate samples per concentration. Samples were cooled and removed from molds. Diameter and height were measured and recorded for each sample. Samples were placed in 1X PBS and stored at 4 **°**C for 24 hours. Diameter and height were measured again for each sample to calculate swelling ratio. Compressive modulus was assessed for each sample using an MTS QT/5, 2N load cell (MTS Systems, Eden Prairie, MN, USA).

### 2.2 Melting and gelation temperature testing

Solutions of 2% w/w LMP and ULMP agarose were prepared in complete DMEM and 1ml of solution was aliquoted into each of 3 separate 2 ml Eppendorf tubes. Gels were cooled to room temperature and allowed to solidify for 2 hours. Tubes containing agarose solutions were placed in a heat block (13259-030, VWR, Radnor, PA, USA) and gently heated. Every 60 seconds, tubes were checked by inversion for flow, indicating melted agarose, and the temperature of the block was recorded when the sample in each tube had melted. The block was heated to 60 °C and turned off. The block was allowed to cool, and tubes were tested every 2 minutes for gelation indicated by lack of flow upon inversion. The temperature at which sample in each tube ceased to flow upon inversion was recorded as the gelation point. This process was repeated 3 times, and the average melting and gelation temperature for each tube was averaged.

### 2.3 PCLS Generation

Mice were euthanized by administration of excess anesthetic (ketamine/xylazine). The trachea was cannulated. Lungs were inflated to 24 mmHg with 2% w/w ULMP agarose in complete DMEM. The trachea was tied with suture and lungs were extracted and placed in conical tubes containing 5 ml complete DMEM on ice for 2hr. Precision-cut lung slices (PCLS) were generated using a vibratome with a 0.076 mm blade (121-4, Ted Pella, Inc., Redding, CA, USA). 1X PBS (10010023, Gibco, Waltham, MA, USA). Individual lobes were separated and allowed to dry. Single lobes were secured to the specimen stage using a thin layer of cyanoacrylate glue, allowing 10 min for curing. The specimen stage was then placed into the PBS bath. PCLS slices were generated at 300 μm thickness, using a frequency of 95 Hz, amplitude of 1 mm, manual interstroke, and velocity of 4 to 6, depending on the stiffness of the tissue. PCLS were placed in complete DMEM media in a 12-well plate (665180, Greiner Bio-One, Monroe, NC, USA) and allowed to rest overnight in an incubator (37°**C**, 5% CO_2_).

### 2.4 PCLS area and agarose digestion measurements

Stitched Montage images of complete PCLS were obtained using a 10x phase-contrast objective on an Agilent Lionheart FX automated microscope (Agilent, Santa Clara, CA, USA). Media was exchanged to complete media containing 100 U/ml agarase enzyme (A6306, Sigma Aldrich). Sample images were obtained as described every 2 hours for the first 8 hours, then every 24 hours for 10 days. PCLS size was quantified by calculating cross-sectional area of each slice over time using ImageJ software (v1.54n, NIH, Bethesda, MD, USA).

### 2.5 PEG Hydrogel Encapsulation

Prior to encapsulation, media was replaced with fresh complete DMEM media containing 100 U/ml agarase. PCLS were encapsulated in polyethylene glycol (PEG) hydrogel in a 6-well ^®^ Tissue Train^™^ stretch plate (TTCF-5001A, FlexCell International, Burlington, NC, USA) immediately following agarase treatment. PEG hydrogels consisted of 2.67 mM (4% w/w) 20 kDa 4-arm PEG-norbornene (A44279-20K, BioPharma PEG, Watertown, MA, USA), crosslinked using 5.33 mM 2 kDa linear PEG-dithiol (SH-PEG-SH-2000, Lysan Bio, Arab, AL, USA) and were photopolymerized with 0.05% w/w photoinitiator Lithium phenyl-2,4,6-trimethylbenzoylphosphinate (LAP) (900889, Sigma-Aldrich) under 365 nm UV light. All hydrogel precursor solutions were made in complete DMEM media.

For PCLS hydrogel encapsulation, 600 μL of PEG precursor solution was added and spread evenly across the bottom of the well and into the surrounding foam ring of FlexCell^®^ Tissue Train^™^ stretch plates, and cured using 365 nm UV light. After polymerization of the base hydrogel layer, 300 μL of 4% w/w PEG was spread evenly on top of the base layer, and the PCLS were placed in the PEG, ensuring it was flat. This layer was then polymerized to encapsulate the PCLS within the hydrogel substrate. For unstretched control samples, PCLS were encapsulated using the same process within standard 6-well paltes.

### 2.6 Stretch Testing

PCLS were stretched using a FlexCell^®^ FX 3000 Tension System (FlexCell International). Tissue Train™ stretch plates containing hydrogels and PCLS were placed on the FlexCell^®^ Tension System in the incubator (37°**C**, 5% CO_2_). An acrylic cover was placed on top of the plates and weighted using 10lb lead weights to ensure a tight seal for the system. Plates that were not being stretched were placed next to the FlexCell^®^ manifold in the incubator. PCLS were stretched at 15% elongation and 10 cycles per minute for 24 hours at a 30% duty cycle.

### 2.7 Bulk RNA Sequencing

After stretching, PCLS were carefully removed from the PEG hydrogel, snap frozen in liquid nitrogen, and stored at -80°C. Samples were sent to the University of Rochester Genomics Research Center for bulk RNA sequencing. Total RNA was isolated using the RNeasy Plus Micro Kit (Qiagen, Valencia, CA). RNA concentration was determined with a NanoDrop 1000 spectrophotometer (NanoDrop, Wilmington, DE) and RNA quality assessed with the Agilent Bioanalyzer 2100 (Agilent, Santa Clara, CA). The quantity and quality of the subsequent cDNA were determined using the Qubit Fluorometer (Life Technologies, Carlsbad, CA) and the Agilent Bioanalyzer 2100 (Agilent, Santa Clara, CA). 150 pg of cDNA was used to generate Illumina-compatible sequencing libraries with the NexteraXT library preparation kit (Illumina, San Diego, CA) per manufacturer’s protocols. The amplified libraries were hybridized to the Illumina flow cell and sequenced using the NovaSeq6000 sequencer (Illumina, San Diego, CA). Single-end reads of 100 nt were generated for each sample and processed to fastq using bcltofastq-2.19.0.

### 2.8 RNA data processing

Sequence files were trimmed for low quality bases using fastp version 0.20.1 with arguments “--length_required 35 --cut_front_window_size 1 --cut_front_mean_quality 13 --cut_front --cut_tail_window_size 1 --cut_tail_mean_quality 13 --cut_tail -y -r.” These were aligned to the mouse genome GRCm38.p6 with Gencode-M25 gene annotation with STAR version 2.7.6a and arguments “--twopassMode Basic --runMode alignReads --outSAMstrandField intronMotif --outFilterIntronMotifs RemoveNoncanonical.” Gene counts were derived using subread version 2.0.1 feature counts with arguments “-s 0 -t exon -g gene_name.” Differential expression was assessed using DESeq2 version 1.34.0 at a false discovery rate of 0.1.

## 3.0 Results and Discussion

Hydrogels of equal weight percentage (2% w/w) formed with ULMP agarose have melting and gelation temperatures 7.27 °C and 6.33 °C lower than those of LMP agarose (**Fig. 1a**). Despite higher compressive modulus of LMP agarose gels across a range of concentrations relative to ULMP agarose (**Fig. 1b**), swelling ratios were not significantly different, and were relatively low, with most measured swelling ratios <1.2 for gels up to 3% w/w (**Fig. 1c**). Hydrogels of >10 kPa have been identified as the minimal compressive modulus for consistent PCLS slicing using a vibratome (data not shown). Swelling ratio could impact the stretch imparted on an inflated lung, as they are typically inflated to physiological pressure, and significant additional swelling would increase the perceived pressure within the tissue. PCLS were generated from mouse lungs inflated with 2% ULMP agarose at 300 μm thickness. Degradation of ULMP agarose from PCLS was achieved using 100 U/ml agarase enzyme within 2hr of treatment and does not impact PCLS viability (**Fig. 1d,e**). On average, a 33.6% reduction in overall PCLS area within 2 hours, with minimal change thereafter through 6 hours (**Fig. 1f**). Interestingly, both agarase treated and untreated PCLS show reduced areas over 7 days in culture, indicating the the reduced melting temperature of the ULMP agarose may be sufficient for removal at 37 °C in a tissue culture incubator. However, this is not to the same extent at observed in agarase-treated PCLS. It is possible that this continued tissue retraction occurs through an alternate mechanism, such as increased mesenchymal cell proliferation or contractility.

**Figure 1.**
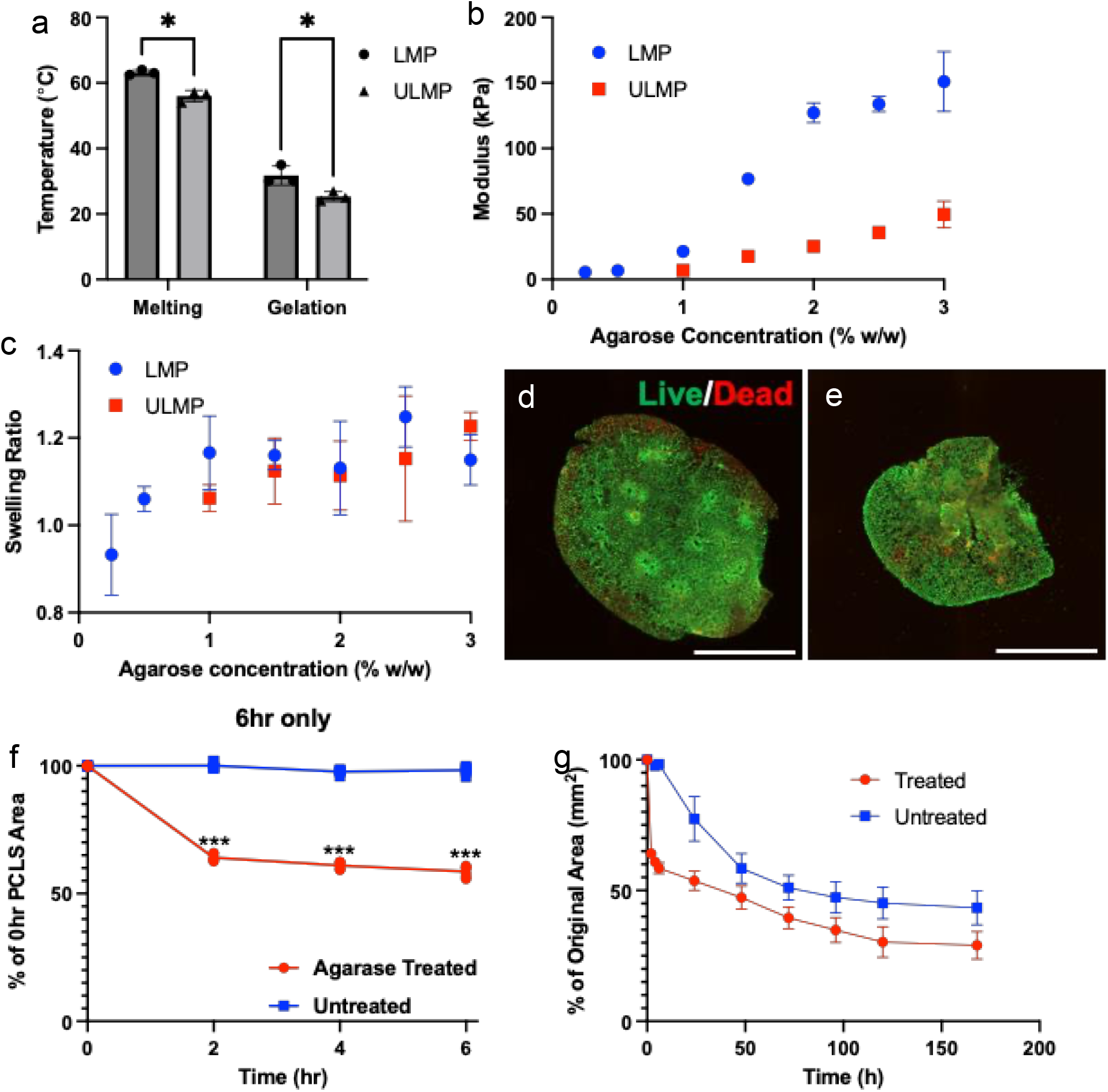
Ultra-low melting point (ULMP) agarose can be used to inflate lung tissues and is removable. A) Melting and geltation temperatures of 2% w/w ULMP and low-melting point (LMP) agarose hydrogels. B) compressive modulus and C) swelling ratio of ULMP and LMP hydrogels up to 3% w/w. Live (green, calcein AM) and dead (red, ethidium homodimer) staining of PCLS derived from ultra-low melt agarose-inflated mouse lung tissue D) before and E) after agarase digestion. Scale bars: 2mm. Plots of PCLS area over F) 6 hrs and G) 168 hours with or without agarose in media. n=3, N=2-3, * p<0.05, *** p<0.001.

Upon removal of agarose, PCLS were successfully encapsulated within PEG hydrogels and incorporated into FlexCell^®^ Tissue Train^™^ stretch plates for cyclic stretch (**Fig. 2a, c**). Application of a measured 15% elongation was measured at an average of 15.01±1.41% upon analysis of stretched and relaxed PCLS (**Fig. 2b, c**). This confirmed that PEG hydrogels were able to transmit stretch from the platform to the PCLS. Critically, after 24 hours of cyclic stretch, PEG hydrogels remained incorporated into the foam anchors of the stretch plates and the PCLS, with no signs of mechanical damage or detachment.

**Figure 2.**
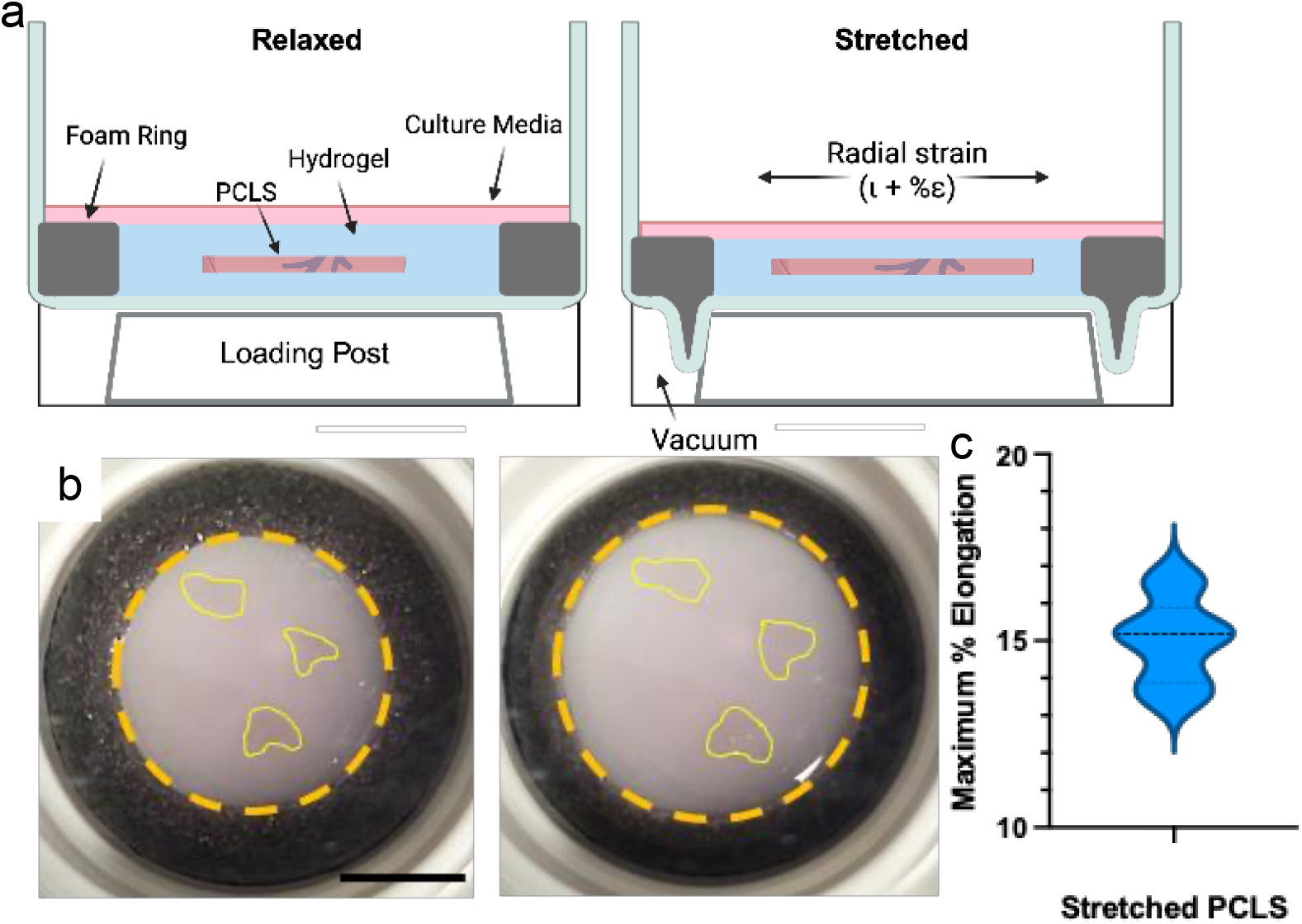
PEG hydrogels allow PCLS incorporation into FlexCell TissueTrain stretch plates without clamping or gluing. A) Schematic of PEG-encapsulated PCLS stretch. B) Image of PCLS in PEG hydrogels integrated into TissueTrain® stretch wells in relaxed (left) and stretched (right) states. Scale bar: 1 cm. Due to tissue translucence, PCLS are outlined in yellow. Hydrogel perimeter is outlined with orange circles. C) Plot of the average % elongation measured by imaging of relaxed and stretched PCLS using a setting of 15% elongation of the FlexCell system. n=9.

Transcriptomic analysis reveals that PCLS exposed to 24 hours of cyclic stretch have distinct gene expression patterns from static samples (**Fig. 3a**). Extracellular matrix gene expression is altered in PCLS that have been stretched (**Fig. 3b**). Extracellular matrix is significantly impacted in mechanically ventilated patients, including neonates and adults.^27-31^ Similarly, genes known to be responsive to stretch also show altered expression patterns in stretched PCLS compared to static controls (**Fig. 3c**). Together, these results indicated that this model may present a viable approach for studying mechanical ventilation-induced injury.

**Figure 3:**
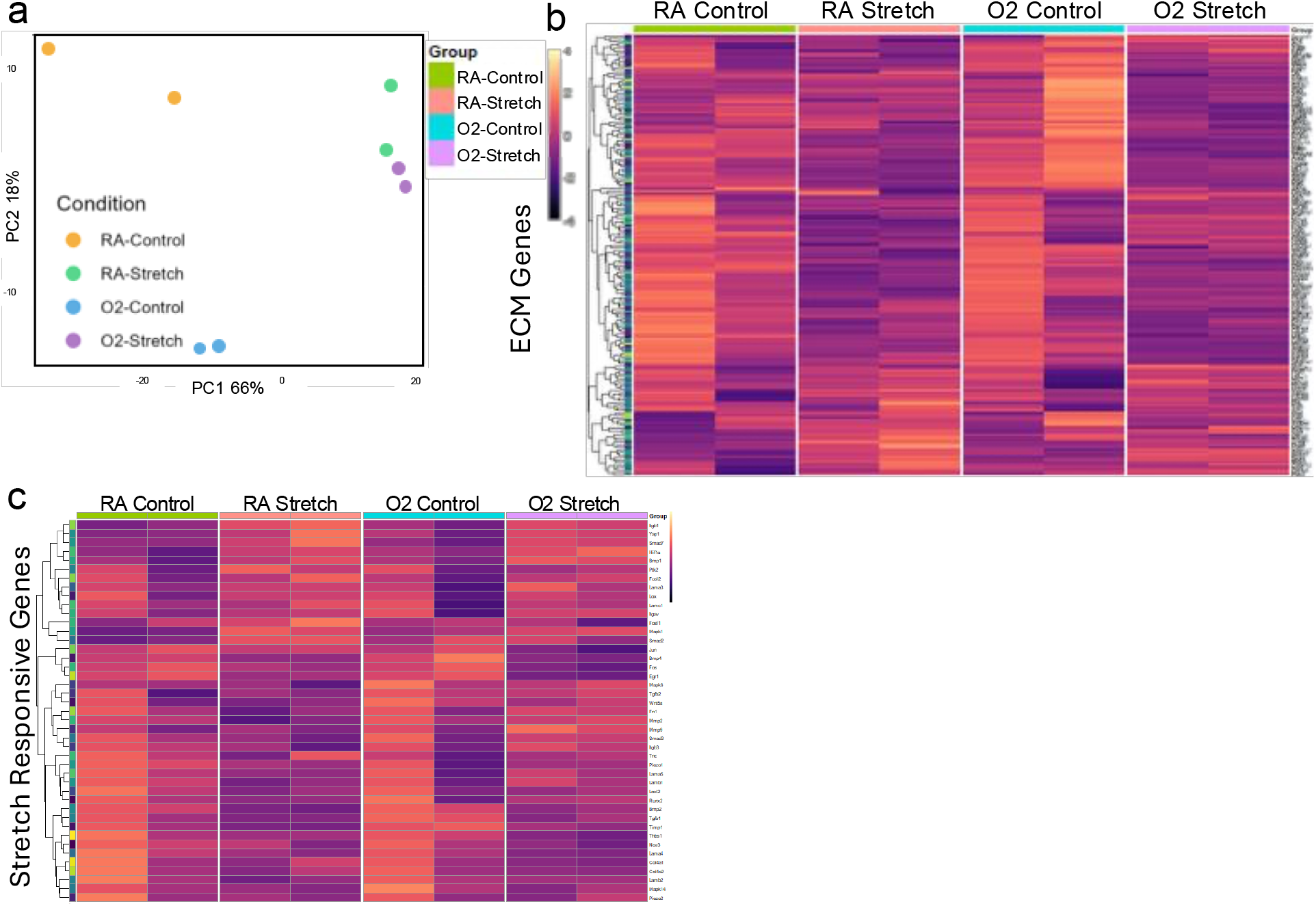
PCLS are stretch-responsive. A) PCA plot of control and stretch samples from adult mice exposed to room air (RA) and 24 hours of hyperoxia before euthanasia (O2). Heat maps of normalized gene expression from RNA sequencing showing B) ECM genes and C) stretch-related genes from matrisomeDB in stretched PCLS and non-stretched controls. N=2, n=3.

## 4.0 Conclusions

These data show an alternate method for lung inflation using ultra-low melting point agarose, which reduces working temperatures and allows for removal by enzymatic digestion. This allows for tissue relaxation and stretch studies to be performed from an unstretched starting state. This material may also improve downstream recovery of nucleic acids and proteins and may improve biochemical analyses. The novel hydrogel-encapsulated cyclic stretch methods apply accurate stretch levels and increase the potential experimental duration to at least 24 hours and do not require damaging attachment methods. This platform has great potential in the ability to use human PCLS, examine the kinetics of mechanical stretch injury and recovery, and can allow for the testing of alternate stretch paradigms, drugs, or the impacts of genetic f factors on stretch response.

## Funding Sources

This work was supported by a University of Rochester University Research Award (to J.A. Mereness). This work was also supported by NHLBI Career Development Award K08HL155491 (to A.M. Dylag).

## Acknowledgements

The authors would like to acknowledge Lananh Ho for help running the FlexCell stretch platform. We would also like to thank the University of Rochester Genomics Research Center (GRC) for facilitating the transcriptomics analyses described in this study.

## Notes

### Competing Interest Statement

The authors have declared no competing interest.

